# Force patterning drives cell flows and 3D spatial order in auditory epithelia

**DOI:** 10.1101/2025.06.26.661757

**Authors:** Julian Weninger, Anubhav Prakash, Sukanya Raman, Raj Ladher, Madan Rao, Karsten Kruse

## Abstract

During development, coordinated cell behavior drives the morphogenesis of epithelia into intricate structures essential for their physiological functions. How this coordination is achieved in epithelia composed of multiple cell types remains unclear. We study development of the avian auditory epithelium comprising sensory hair cells (HCs) and non-sensory supporting cells (SCs). Initially, HCs and SCs are arranged into mosaics by Notch-Delta signaling. During development, HCs partially extrude from the epithelium, establish a ten-fold gradient in apical surface area across the tissue, and intercalate with SCs to form near-hexagonal order. Using a combination of experiments and a 3D-vertex model, we show that an increase of contractility at the apical junctions between SCs compared to those between HCs and SCs drives spatial order within the epithelial plane and along the apical-basal axis. A faster increase of HC apical area at one end of the epithelium than the other leads to opposite fluxes of HCs and SCs and establishes the observed gradients in apical surface area and density of HCs along the auditory epithelium while maintaining a uniform degree of hexagonal order throughout. Our findings reveal that patterned junctional contractility can spatially coordinate cell behavior across both the plane and depth of a mixed epithelium, providing a general mechanism for generating complex three-dimensional tissue architectures during organogenesis.

## I. INTRODUCTION

Precise spatial organization of cells into multicellular arrays is critical for the functional architecture of organs. Such patterning occurs not only at the luminal surface but also across the luminal-lateral plane, for example, the planar-polarized orientation of hairs in the Drosophila wing [1, 2] or the stratified positioning of hair follicles in the skin [3]. During development, luminal and lateral organization are often established simultaneously, yet in a coordinated manner, to sculpt tissues into their mature forms. Despite their importance, the mechanisms that orchestrate this concerted spatial organization remain poorly understood. To address this, we investigated the developing chick auditory epithelium, known as the Basilar Papilla (BP), which exhibits both apical surface patterning and lateral intercalation of distinct cell types, sensory hair cells and non-sensory supporting cells, making it an ideal system to uncover how planar and vertical organization emerge together during epithelial morphogenesis.

In the apical plane, the auditory epithelium presents a fascinating pattern of sensory hair cells (HCs) and non-sensory supporting cells (SCs), where HCs organize into a near hexagonal arrangement while being intercalated by SCs [4, 5]. Along the apical-basal axis, HCs and SCs are connected apically, and only SCs extend to the basal lamina [6, 7], making the tissue quasi-stratified. Additionally, the auditory epithelium is tonotopic along the proximal-distal axis, in which HC shapes vary strongly among others [8]. This shape variation includes the HC bundles, the apical protrusions that mechano-transduce acoustic stimuli, the HC apical surface, and the lateral arrangement of HCs with respect to the SCs. In particular the shape of the HC bundles is critical, as it contributes to the increased sensitivity of the organ to specific frequencies, where longer protrusions are more sensitive to the longer wavelengths of lower frequencies [9]. How these spatial variations are regulated and mechanically coupled is largely unknown.

Mechanical forces play an important role in shaping, dividing, and rearranging cells within the epithelial plane as well as along the apical-basal axis during development. For example, in the fruit fly *Drosophila melanogaster*, mesoderm invagination results from localized contractions of the actomyosin network [10], and neural tube closure in chick embryos relies on actomyosin accumulation at specifically oriented cell-cell junctions [11]. Activation of actomyosin contractility can be biochemically regulated, and this regulation is in turn coupled to tissue mechanics, geometry, and material properties. The regulatory role of mechanics was found to be crucial for wing development [12–14] and germ band extension [15, 16] in *D. melanogaster*.

In tissues composed of multiple cell types, like auditory epithelia, mechanical and biochemical properties can vary between cells of different types and between junctions formed by different cell types. These heterogeneities in chemo-mechanical properties of cells and junctions drive the organization of cells [5, 17–19] and segregate germ layers during development [20, 21]. During the development of the BP, lateral inhibition via Notch-Delta pathway picks delta high HCs, surrounded by SCs. Once differentiated, the apical junctions between two SCs are enriched with di-phosphorylated form of the regulatory light chain (ppRLC) of non-muscle Myosin II, leading to higher junctional contractility and driving cellular reorganization [5]. However, the role of heterogeneous chemo-mechanical properties of HCs and SCs in three-dimensional organization remains elusive.

In this study, we combine experimental and theoretical approaches to uncover how spatial differences in junctional contractility drive three-dimensional cell rearrangements during basilar papilla (BP) development. These rearrangements are sufficient to generate both quasi-stratification and the positional ordering of hair cells (HCs). We show that gradients in tissue mechanics produce relative movements between HCs and supporting cells (SCs), leading to the emergence of the graded surface area of HCs and thereby contributing to BP tonotopy. Together, our findings highlight junctional force patterning as a key mechanism underlying the coordinated morphogenesis of the BP.

## II. RESULTS

### A. BP displays uniform spatial order despite 10-fold differences in HC apical area

To investigate the development of spatial order across epithelia, we immunostained BP with F-actin to mark cell boundaries and Hair cell antigen (HCA) to identify HCs. At embryonic day (E) 8, when the HCs are first apparent, the apical surface area of HCs are similar across the proximal-distal axis of BP. By E14, as the BP matures, and similar to previous observations, we found a 10-fold gradient of HCs apical surface area along the P-D axis. Despite this strong gradient, HCs locally showed a positional order in both chick BP and mouse auditory epithelium. To investigate the tissue-wide angular regularity of HC positions along the P-D gradients of BP, we use the generic *p*-atic order parameter *ψ*_p_ with *p* = 3, 4, 5, … that measures degree of discrete rotational symmetries, including hexatic symmetry for *p* = 6 (SI Appendix, Sect. C 2 a). We find that the hexatic phase emerges throughout the BP from an originally unordered tissue, with *ψ*_6_ *>* 0.5 at E14 (Fig. 1e, SI Fig. S1a-c). Although we did not find a long range order probably because of a gradient in the apical surface area, the hexatic order across BP suggested the generation of a positional order across the BP. In previous work, we found that local hexatic order of BP is driven by HC apical area and is maximal for a precise value of this area [5]. Hence, we decided to investigate the underlying mechanism that couples HC area with the hexatic order.

**FIG. 1.**
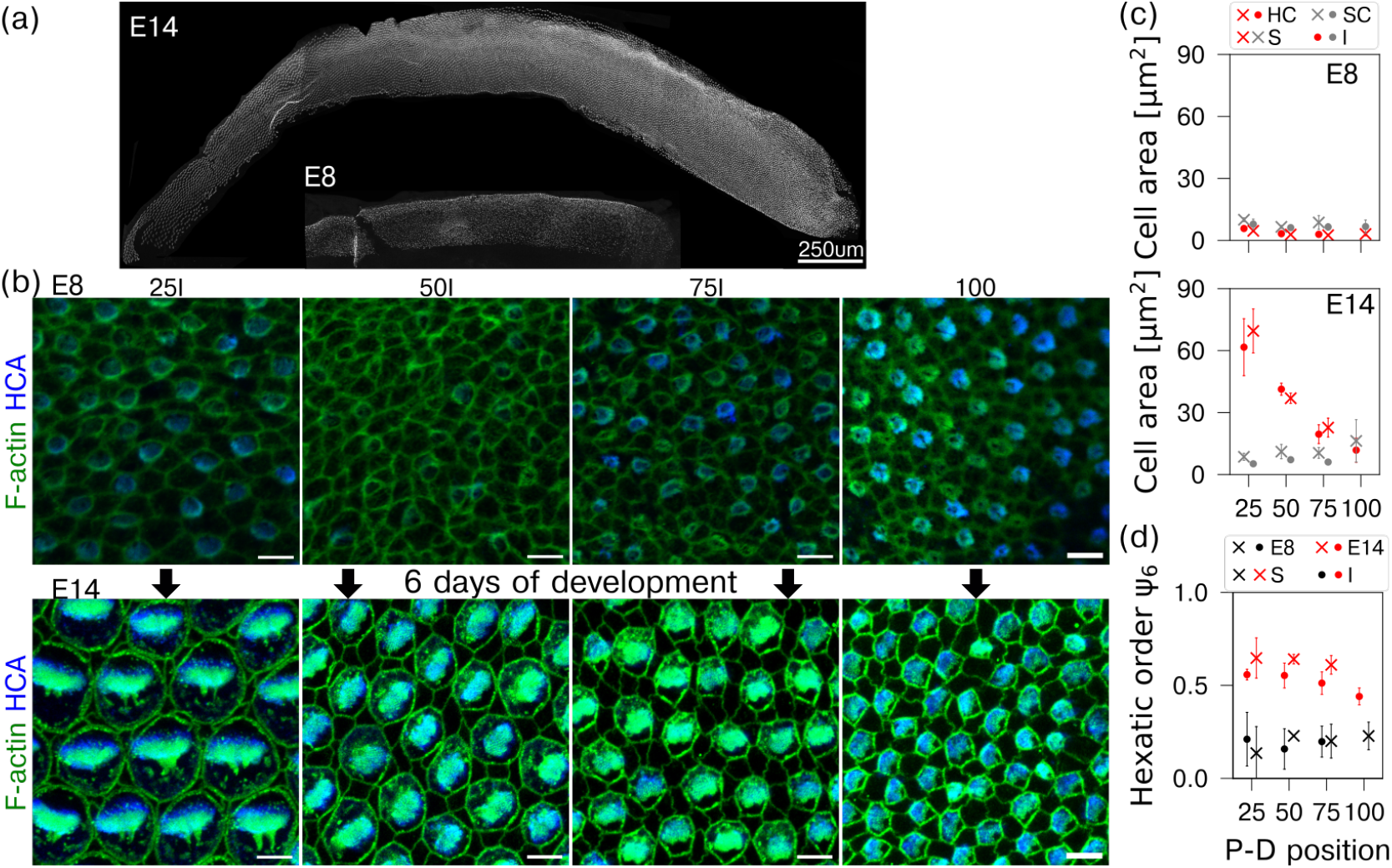
Positional order develops in basilar papilla (BP) despite gradients in apical surface area. a) HC area along the BP at embryonic day (E)8 and E14. Inset: Hair cell antigen (HCA) staining (right) and segmentation (left). Scale bar 100 *µ*m (inset 10 *µ*m). b) HC arrangement along the proximal (P) – distal (D) axis (25 %, 50 %, 75 %, and 100 %) in the inferior (I) part of BP at E8 and E14. Green: F-actin, blue: HCA. Scale bar, 5 *µ*m. c-d) Cell area (c) and hexatic order parameter 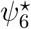 (d) along the inferior (I, circles) and superior part (S, crosses) of BP at E8 and E14 for HCs (red) and SCs (gray). E8 is shown in black in (d). Crosses: means of sample means, error bars: standard deviation of means. Symbols in (c,d): sample means.

### B. HC apical area growth is accompanied by quasi-stratification of BP

In mouse auditory epithelia, the change in apical surface area of HCs is driven by growth and additional shape changes [6]. To measure cell volume and shape changes in BP, we sectioned the BP into 20 *µ*m thick slices and stained for Myosin 7a to mark the cytoplasm of HCs, DAPI to mark the nuclei of both HCs and SCs, and Phalloidin to mark cell apices (Fig. 2a). We obtained fluorescence images by confocal microscopy of individual cross-sections and reconstructed 3D images. In these images, we segmented HC volumes based on the Myosin 7a staining and nuclear volumes based on the DAPI staining. HC volume increased significantly between E8 and E10 from (150 ± 30) *µ*m^3^ to (200 ± 50) *µ*m^3^ (mean ± std. dev.), but not between E10 and E14 (less than 25 %, Fig. 2b). Throughout development, proximal HCs had smaller volumes than distal HCs (Fig. 2b). The gradient in volume is opposite to the one in apical surface area. Thus, the increase in volume cannot account for the up to 10-fold decrease in apical area along the P-D axis (Fig. 1c).

**FIG. 2.**
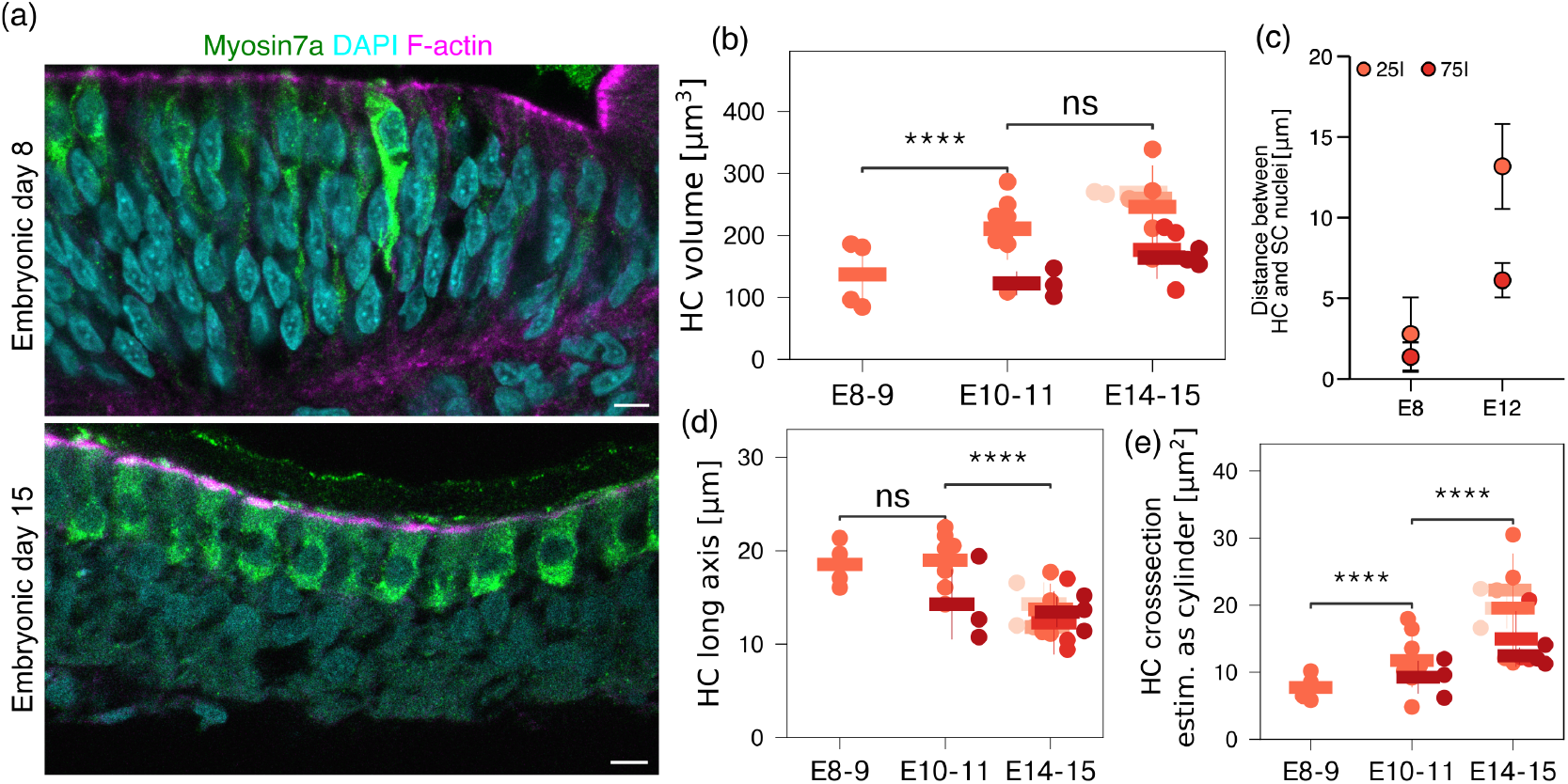
HC area is controlled by quasi-stratification of HCs and SCs along the apical-basal axis. a) Cross-sections of BP at E8 and E15 stained for Myosin 7a (green, HCs), DAPI (cyan, HC and SC nuclei), and F-actin (magenta, apical plane of cells), showing the apical location of HCs and their nuclei compared to SCs. b-e) Measurements of HC volume (b), distance between HC and SC nuclei (c), HC height (d), and HC cross-section area at different stages. Dots: tissue means, horizontal bars: mean of means, error bars: standard deviation of means. n.s.: non-significant (*p >* 0.05), ***: *p <* 0.001, ****: *p <* 0.0001, Man-Whitney-test. Scale bars: 5 *µ*m (a), 1 *µ*m (f).

We further investigated the apical-basal structure of the BP. While HC and SC nuclei are found at similar positions along the apical-basal axis at E8, their vertical distance increases significantly until E15 (Fig. 2c). This apical-basal segregation is furthermore apparent in the 3D remodeling of HCs: At E8 the HC, as labeled by Myosin 7a, lie on the basal membrane, however by E15 they retract apically (Fig. 2a, S2a). This quasi-stratification results in an almost 2-fold decrease in HC apical-basal height (Fig. 2d-e) and is complemented by an hourglass-like constriction of the HCs below the apical plane (Fig. 2a) further increasing HC apical area.

### C. Junctional force pattering in the lateral plane drives quasi-stratification

To understand what is driving this quasi-stratification, we turn to tissue mechanics. Our segmentation shows that HCs are separated from each other (Figs. 1b, 2a). In previous work, we identified differences between the contractility of HC-SC and SC-SC junctions to drive HC separation in the apical plane [5]. SC-SC junctions are enriched in double-phosphorylated Regulatory Light Chain (pp-RLC), a component of the non-muscle Myosin-II motor hexamer, which governs their increased contractility compared to HC-SC junctions. A super-resolution microscopy of junctions showed the pp-RLC is present at the apical plane (Fig. 3a, SI Movie 1), suggesting a higher contractility at apical junctions between two SCs. To understand the role of increased apical contractility in SC-SC junctions for the 3-dimensional reorganization of BP, we turn to a physical description of BP mechanics in the cross-sectional plane. We consider a minimal SC-HC-SC arrangement, where all cells are polygons of equal height and area (Fig. 3b). The state of the system is determined by the minimum of the effective energy

**FIG. 3.**
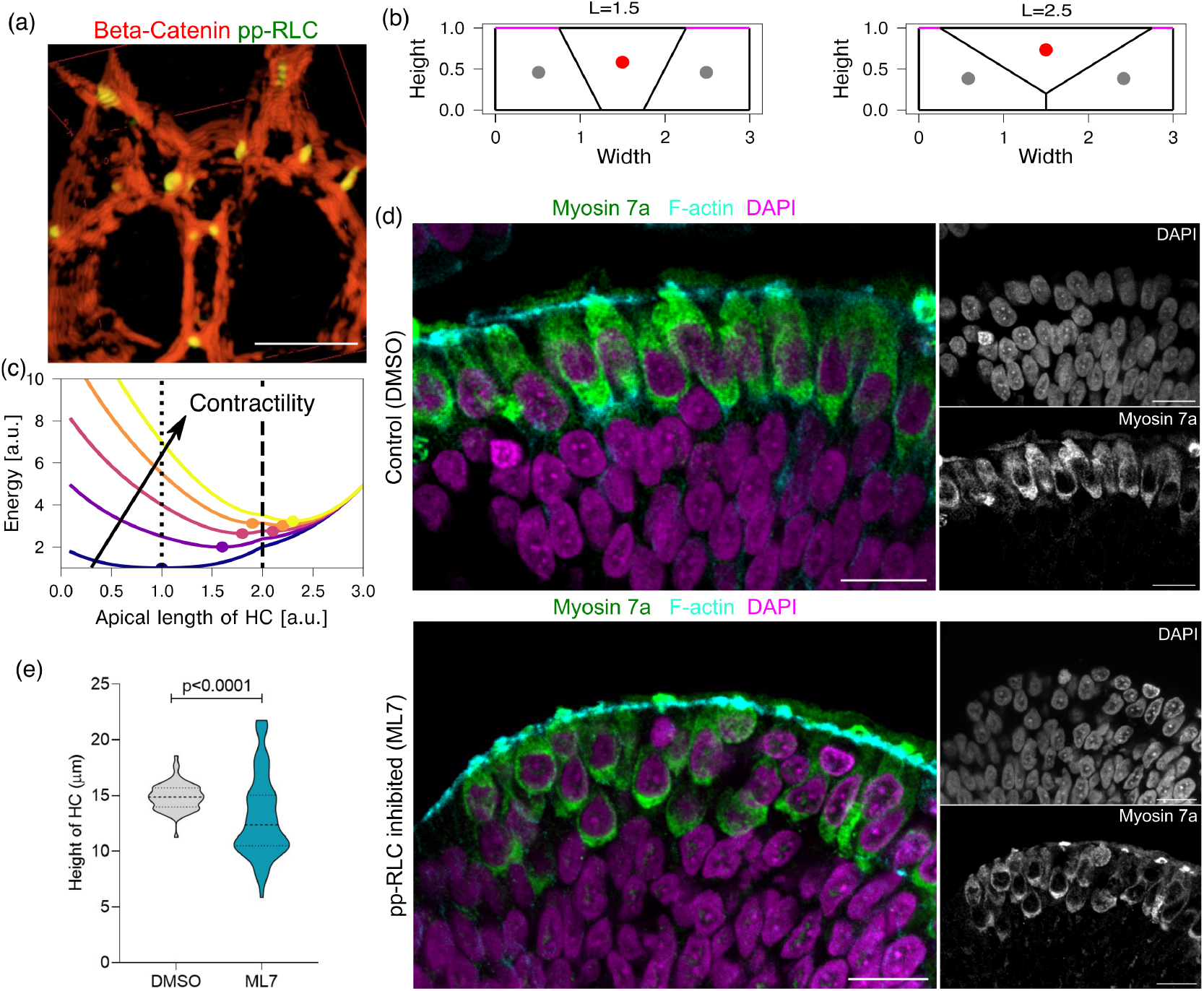
Quasi-stratification is controlled by pp-RLC through apical junctional contractility. a) Super resolution image from structured illumination microscope showing *β*-catenin (red) and two bands of pp-RLC (green, white arrows) at the apical junctions between two SCs at E10. b) Schematic of lateral configurations of a HC (red dot) between two SCs (gray dots) for different apical HC lengths *L*. SC junctions with increased contractility Γ are indicated in magenta. HCs detach from the base at *L* = 2. c) Energies of SC-HC-SC configurations as a function of *L* for different values of Γ. Dots: energy minima. d) Cross-sections of BP at E10 treated with DMSO (Control, upper) and ML7 (lower) stained for Myosin 7a (green, HCs), DAPI (magenta, HC and SC nuclei), and F-actin (cyan, apical plane of cells). e) Distribution of HC heights in DMSO and ML7 treated samples. Data from 3 tissues each, N=69/72 DMSO/ML7, resp.; horizontal lines indicate mean and std. dev.; *p <* 0.0001, student t-test. Scale bars: 5 *µ*m (c).

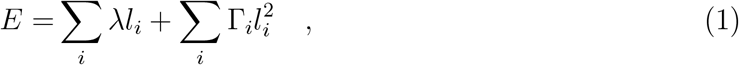

where the index *i* labels junctions and *l*_i_ denotes their lengths. The first term in Eq. 1 represents the tension of junctions, whereas the second term accounts for junctional contractility [22]. For simplicity, we set the line-tension parameter *λ* to be 1 on all junctions. Based on the distribution of pp-RLC, we consider the contractility parameter Γ_i_ to be non-zero and equal to Γ only on apical junctions of SCs (Fig. 3b,c). The contractility parameter Γ is the only free parameter in this description.

Considering a single mode of deformation, such that the height *h* and the area *A* of all cells are constant and equal to 1, we plot the effective energy as a function of the apical HC length *L* (Fig. 3c). This displays a single minimum at the columnar configuration *L* = 1 for low values of the contractility Γ. With increasing values of Γ, this length increases. There is a critical value of Γ at which a second minimum appears at *L* ≥ 2 corresponding to basal detachment of the HC (Fig. 3c). For Γ ⪆ 1.6, this second minimum becomes the global minimum. Eventually, the primary minimum disappears and the detached state becomes the unique minimum. This indicates that patterned apical junctional contractility can alter the apico-basal arrangement of cells and induce a quasi-stratification in an alternating arrangement of HCs and SCs. In agreement with our experimental observations, our analytical results show that the increased apical contractility of SC-SC junctions results in a reshaping of HCs, where their apical area increases and apico-basal height decreases. This leads to basal detachment of HCs and potentially stratification of HC and SC nuclei (Fig. 2a-c).

To test this experimentally, we used an ex-vivo organ culture of BP in presence of pharmacological inhibitors of myosin light chain kinase, ML7. In the presence of ML7, the distance between the HC nuclei and SC nuclei was reduced compared to the control samples (Fig. 3d). Even though the average HC height decreases (Fig. 3d-e), in contrast to our theoretical predictions, the increase in variability of HC height and vertical distance between HC nuclei (Fig. 3d-e) suggest a loss in vertical organization of BP. Our theoretical model does not account for variability in nuclei positioning and influences on the HC shape. This reduction in vertical organization suggests the presence of pp-RLC at the SC-SC junctions contributes to the quasi-stratification of BP. Previous studies have shown the presence of pp-RLC driven higher contractility on the SC-SC junctions is critical for the development of hexagonal arrangement of HCs at the apical plane, suggesting a mechanistic that drives both in-plane organization and quasi-stratification.

### D. Quasi-stratification is coupled to ordering transition

To gain insights into the this mechanism, we turn to a 3D vertex model, where the positions of the vertices in the third dimension are appropriately constrained to reflect the BP geometry.

Vertex models for tissues are determined by a work function *W* that associates an energy with a given configuration of the cells that are represented as polygons or polyhedra [23]. In 3D, this work function depends on the cell volumes *V*_α_ and cell-cell interfaces *I*_⟨α,β⟩_, where *α* indexes a cell and ⟨*α, β*⟩ denotes a pair of neighbouring cells [24]. Explicitly, we write

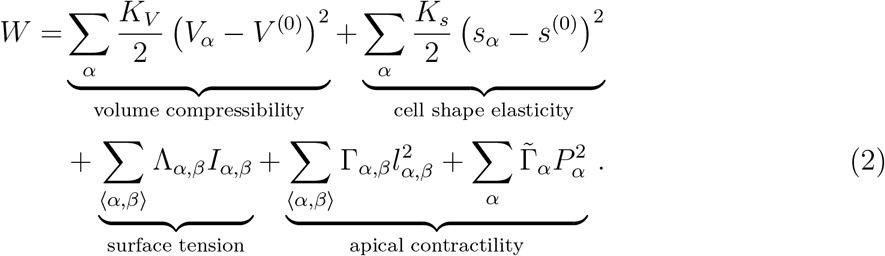

This description penalizes deviations from a target cell volume *V* ^(0)^ and a target cell shape *s*^(0)^ by the elastic constants *K*_V_ and *K*_s_, respectively. The cell shape is given by the normalization of cell surface area *S*_α_ to the target cell volume *s*_α_ = *S*_α_ (*V* ^(0)^)^−2/3^. Interfacial tension and junctional contractility are characterized by the parameters Λ_α,β_ and Γ_α,β_, respectively. Note that we locate the contractile features of the junctions at the apical side of the tissue, corresponding to the location of the Myosin-II complex in tight junction (Fig. 3a, SI Movie 1). Therefore, the contractility is expressed in terms of the junctional lengths *l*_α,β_ in the apical plane (Fig. 4a). Finally, we also consider apical contractility of cells, characterized by the parameter 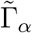 and the cell perimeter *P*_α_.

**FIG. 4.**
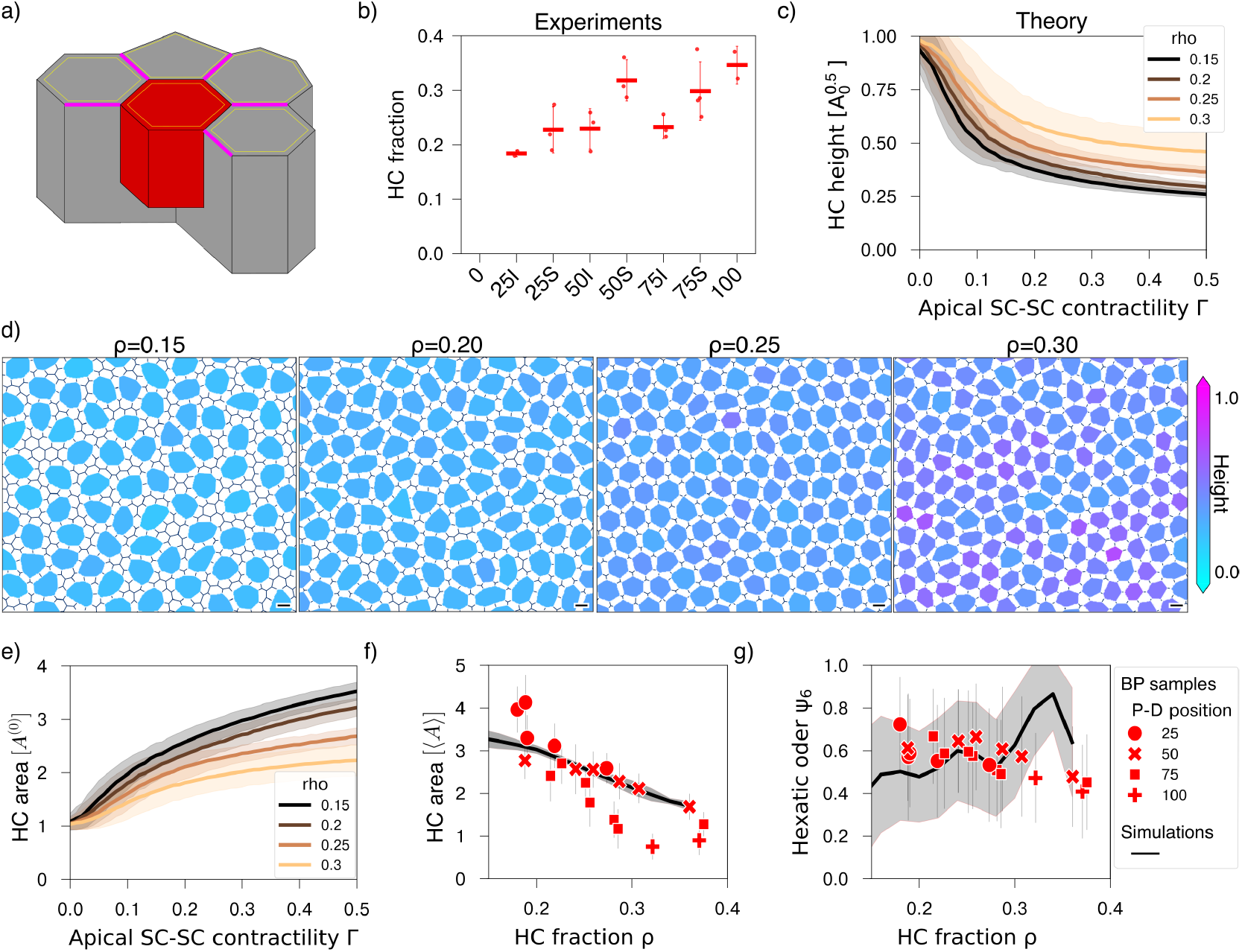
Quasi-stratification drives positional ordering and compensates differences in HC fraction along the BP. a) Sketch of the 3D vertex model, where HCs (red) can differ in height from surrounding SCs (gray). The volume below the HC is distributed to the neighbouring cells. Magenta lines represent increased apical SC-SC junctional contractility. b) Variation in HC fraction *ρ* across the BP at E14. c) HC height in the 3D vertex model as a function of SC-SC contractility Γ for different values of *ρ*. d) Apical patterning of simulated BP at steady state for *ρ* = 0.16, 0.20, 0.26, and 0.32. HC height is color coded. e) HC area as a function of apical SC-SC contractility for different values of *ρ*. f, g) HC apical area (f) and hexatic order (g) in BP samples (symbols) and the 3D vertex model (line) as a function of *ρ*. In BP, HC area was measured in units of the mean cell area. Lines and points: mean values, shaded areas and crosses: standard deviation. BP data points from individual images.

In the following, we restrain 3D geometries allowed in the vertex model: We start our simulations with a columnar arrangement, where all cells are described as prisms with identical apical and basal polygonal shapes and height *H*^(0)^. During minimization of the work function *W*, HCs are allowed to vary their height, *H*_α_ ≤ *H*^(0)^, as shown in Fig. 4a. Explicitly, we model the positions of vertices in the flat apical plane and basal vertices are a projection of those apical vertices to a z-position, given by the cell’s height. As a consequence, the interfaces with neighboring cells remain vertical. The space beneath an HC with *H*_α_ *< H*^(0)^ is filled by the neighboring SC (SI Appendix, Sec. 1).

To minimize the work function, we use a gradient descent method [25]. We initiate the minimization process with a random distribution of columnar cells that all have the same values 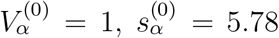 and *H*_α_ = *H*^(0)^ = 1. We assign a fraction *ρ* of the cells to be HCs, whereas the others are SCs. In the BP at E14, we measured the HC fraction to vary between 15 % and 30 % from the proximal to the distal end (Fig. 4b). Based on the distribution of pp-RLC, we consider increased junctional contractility on SC-SC junctions, Γ_S,S_ *>* 0, whereas Γ_H,S_ = Γ_H,H_ = 0. All other parameters are independent of the cell or the cell-cell junction types (SI Appendix, Table 1). To facilitate cell-neighbor exchange, we allow the surface tension Λ_α,β_ to fluctuate.

In the absence of apical junctional contractility at SC-SC interfaces, Γ_S,S_ = 0, the columnar configuration of HCs is stable (Fig. 4c). With increasing Γ_S,S_, we observe 3D remodeling of HCs. HC height decreases with contractility (Fig. 4c), which is similar to the results reported in Fig. 3d,e. Furthermore, independently of the initial condition, HC-HC contacts are essentially absent in steady state [5]. A decrease in HC height implies an increase of HC apical area with *A*_α_ ≈ *V/H*_α_ with *V* ≈ *V* ^(0)^ (Fig. 4d,e). We observe that the decrease in HC height *H* depends on the HC fraction *ρ* and is larger for larger values of *ρ*. This could be due to reduced overall contractility as the fraction of SC-SC junctions decreases or due to increased proximity to other HCs with increasing *ρ*. The saturation of ⟨*H*_α_⟩ at high SC-SC contractility, Γ_S,S_ ≳ 0.3, suggests that the second mechanism dominates (Fig. 4c).

To test the second mechanism, we compared simulations at Γ_S,S_ = 0.4 to individual images taken at E14 across the BP. As a function of the HC fraction *ρ*, we find good agreement between experiment and simulations where the relative apical area of HCs and SCs ⟨*A*⟩_HC_*/*⟨*A*⟩_SC_ decreases with increasing *ρ* (correlation coefficient *x* = 0.85, Fig. 4f). Similarly, the hexatic order parameter *ψ*_6_ varies around 0.6 in both, experiments and simulations, although in simulations the hexatic order increases as *ρ* approaches 1/3, whereas the experimental data show a slight decrease with *ρ* (Fig. 4g). In the absence of HC-HC contacts, there is a mechanical limit to the HC area increase. When this dense packing is reached, the hexatic order is maximal and HCs cannot extrude further. This observation suggests that hexatic order is a consequence of SC-SC contractility that balances HC area increase through 3D remodeling and effective HC-HC repulsion; thus hexatic order depends on HC area and HC fraction.

### E. Force patterning drives HC separation dependent on tissue rheology

During development, the dynamics of Notch-Delta signaling distinguishes Delta-high HCs from surrounding Notch-high SCs through lateral inhibition [26, 27]. This biochemical signaling explains the initial spatial separation of HCs at E8. However, it is unclear how HC separation can be maintained during the subsequent tissue scale convergent extension between E8 and E14 (Fig. S1a). Indeed, convergence-extension of tissues is often accompanied by major cell rearrangements [12–14]. One might expect these rearrangements to lead to numerous HC-HC contacts. However, during the whole developmental process, HC-HC contacts are rare in the BP [5, 28].

To address this, we need to take into account the rheology of the tissue over the timescale of BP convergence-extension and positional ordering between E8 and E14. Hence, we study our 3D vertex model in the presence of an imposed shear (Fig. S3e) (SI Appendix, Methods). Duclut et al. [29] show that the homogeneous 2D vertex model shows linear viscous response at high surface-tension fluctuation amplitudes and shear-thinning at low fluctuation amplitudes.

Applying a shear deformation in the absence of junctional heterogeneity Γ_S,S_ = 0, we observe that many HCs contact other HCs with an average 1 HC contact per HC at steady state. However, HC-HC contacts are reduced in the presence of contractile SC-SC junctions, Γ_S,S_ *>* 0 (Fig. S3e,f). This reduction is partial for small fluctuation amplitudes, where shear thinning promotes the formation of HC-HC contacts. In contrast, sufficiently large contractility of SC-SC junctions resolves essentially all HC-HC contacts. The average number of HC-HC contacts is non-monotonic as a function of the fluctuation amplitude, because, for small values of Γ_S,S_ *>* 0, they are even more frequent than in the absence of SC-SC junction contractility (Fig. S3f). This increase in HC-HC contact number shows that, for small fluctuation amplitudes, the SC-SC contractility is insufficient to separate HCs.

The formation of HC-HC contacts is closely related to cell elongation. The latter is proportional to residual shear stress [29] and exhibits a minimum for intermediate fluctuation amplitudes (Fig. S3g). This minimum coincides with the minimum of HC-HC contacts. Note that the corresponding fluctuation amplitude depends on the value of SC-SC contractility. This is because the tissue becomes stiffer with increasing contractility (Fig. S3e). We also note that the cellular diffusion constant *D* ≈ 1 × 10^−3^ V_0_*/*(H^(0)^*τ*_Λ_) in simulations with zero shear rate and for fluctuation amplitudes that yield minimal HC-HC contacts (Fig. S3h). Together, fluidity of the tissue and a repulsive force between HCs mediated by SC-SC contractility suppresses HC-HC contacts.

Based on the results from our vertex model simulations, we infer that the developing BP, for which cell elongation is small compared to the convergent-extension of the BP and from which HC-HC contacts are absent, is viscous rather than elastic over the timescale of 6 days of development. We, therefore, choose in all our simulations rheological conditions that are close to the conditions where HC-HC contacts vanish by choosing an appropriate fluctuation amplitude.

### F. Spatial gradients induce cell flows

Our results in the previous section show that the hexatic order displayed by HCs in the BP results from an increase in HC area, which is dependent on HC fraction. As noted above, HC area and fraction display opposite gradients along the proximal-distal axis (Fig. 1c and 4b). In the following, we discuss how HC fraction is regulated and show that it too is governed by tissue mechanics.

Until E10, while HCs are differentiating, a salt-and-pepper arrangement of HCs and SCs with hardly any HC-HC contacts emerges through Delta-Notch interactions [30]. Furthermore, HCs and SCs have similar apical surface areas, and both cell types are columnar. After E10, cell differentiation slows down and cell proliferation essentially stops [5]. Hence, most rearrangements are due to cell intercalations and quasi-stratification of BP. To investigate a possible origin of HC rearrangements, we use our 3D vertex model.

Our experiments show that HC area growth speed decreases along the P-D axis of the BP (Fig. 1c,d). We incorporate this observation by varying 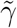 in 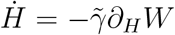, where *H* is the height of a HC (Fig. 4a). This corresponds to a variation of the dissipation rate associated with HC extrusion. Explicitly, we vary 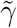 by a factor of 5 along the P-D axis, 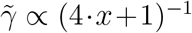, where *x* is the distance from the proximal end. Alternatively, we could also apply a gradient in the contractility Γ as we discuss in SI (Fig. S5).

In the presence of a gradient in HC extrusion speed, HC area increases 3-fold in proximal positions, where HC extrusion is faster, while SC areas decrease 2-fold (Movie 2, Fig. 5a). As HC areas increase, HC-HC distances decrease. Due to SC-SC contractility, HC-HC contacts remain rare implying an effective repulsion force between HCs. In combination, this generates a HC flux towards the distal end (Fig. 5b). At the distal end, HC area changes more slowly and remains smaller with larger HC-HC distances. Accordingly, HCs can accumulate following their displacement from the proximal end. At steady state, we observe a 3-fold gradient in HC area between the proximal and the distal ends (Fig. 5a,c). Concomitantly, an opposite gradient in HC fraction is established as a result of the cell displacements (Fig. 5d). HC fraction varies between 0.2 and 0.3 covering the same range as in BP.

**FIG. 5.**
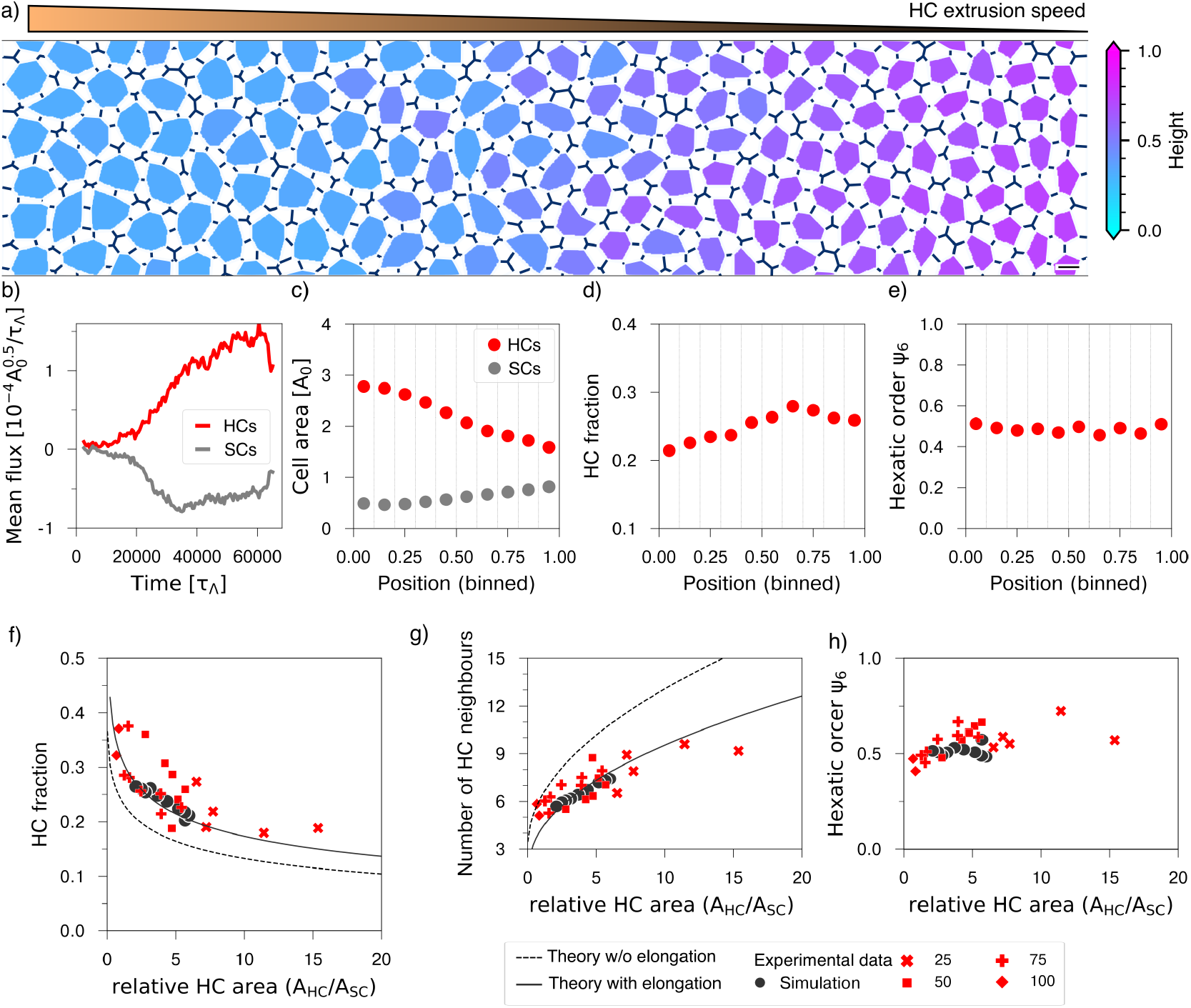
Proximal-distal gradient in the HC extrusion speed drives cell flows and establishes hexatic order across the BP. a) Steady state of a 3D vertex-model simulation with graded HC extrusion speed 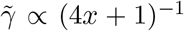 along the P-D axis position *x*. b) Mean cell fluxes of HCs (red) and SCs (gray), respectively, oriented towards the distal and proximal end. Fluxes have been calculated over 5000 *τ*_Λ_ time intervals. c-e) HC and SC area (c), HC fraction (d), and hexatic order (e) along the horizontal axis of the simulation at time 60 000 *τ*_Λ_. f-h) HC neighbor number (f), HC fraction (g), and hexatic order (h) as a function of relative HC to SC area from simulations (red) and BP tissues (black) at E14. Analytical estimate of the number of HC neighbors (f) and HC fraction (g) without (dashed line) and with SC elongation (solid line) (SI Appendix, Sect. C 4). Data points indicate sample means, horizontal bars means of means and error bars standard deviations of means. P-value from Mann-Whitney U-test comparing E10 and E14 means.

Throughout the simulated tissue, we find uniform hexatic order with *ψ*_6_ ≈ 0.5 (Fig. 5e). For example, at *x* = 0.15 ± 0.05, we find a HC fraction *ρ* ≈ 0.2 and HC area ⟨*A*⟩_HC_ ≈ 3*A*_0_.

This is in agreement with our previous results (Fig. 3d,g). Similarly, for *x* = 0.85 ± 0.05, we find a HC fraction *ρ* ≈ 0.3 and HC area ⟨*A*⟩_HC_ ≈ 2*A*_0_. In contrast to the results presented in Fig. 4, here, the HC fraction was not imposed but emerges as a consequence of the HC flux. We conclude that the faster HC extrusion in the proximal part establishes an HC area gradient that drives a cell flow across the BP. The increase in HC fraction mechanically limits the distal HC area increase.

In the Appendix, we provide an analytical estimate of the HC fraction using a geometrical construction. We consider two scenarios, one without SC elongation and another with SC elongation *q*. We observe that the first scenario provides an upper limit to our experimental and numerical data (Fig. 5f). We thus fit the SC elongation to the data from BP at E14 in Figure 5f, obtaining *q* = 0.129(14). In BP, a direct measurement of SC elongation at different positions reveals 0.2 ⪅ *q <* 0.6 (Fig. S4e). Our analytical estimate with SC elongation recapitulates well the HC fraction obtained from individual images of different BP positions at E14 (Fig. 5g). We also find a good agreement between our estimate and our simulated tissues regarding HC neighbor number and HC fraction (Fig. 5f,g). Furthermore, hexatic order in simulated BP is similar to E14 BP (Fig. 5h).

Since it is not possible to live-image the entire BP for several days and directly measure cell fluxes, we analyzed the HC fraction along the tissue for several fixed BP at different stages between E8 and E14 (Fig. S4g-j). The total number of HCs increased between E8 and E10 after which it remained constant until E14 (Fig. S4h). Locally, we observed changes throughout the whole developmental process. At proximal positions, 25S / 25I, the HC fraction *ρ* increased between E8 and E10, reaching a fraction *ρ* = 0.24 ± 0.02 at E10 (Fig. 4i, S4f). At this stage, no tissue displayed *ρ <*22 %. By E14, *ρ* decreased again proximally with some tissues having *ρ* ≈18 %. This is in contrast to the distal region, where *ρ* increased on average from E8 to E14 reaching up to 38 % (Fig. S4g). For a finer spatial resolution, we observed a decrease of HC number in proximal regions from E12 to E14, whereas it increased simultaneously in distal regions (Fig. S4i,j). These changes in HC distribution are compatible with a cell flux, but could in principle also have contributions from transdifferentiation of HCs into SCs.

## III. DISCUSSION

In this paper, we show that the development of cellular organization of the BP’s luminal and lateral surfaces are geometrically and mechanically coupled. Our work shows that a purely 2D description of tissue organization is inadequate to describe development of the mature BP. Instead, one needs to account for three-dimensional cellular deformations, in particular, the increase in HC apical surface area and concomitant decrease in HC height, which leads to a quasi-stratification of the epithelium. Using a 3D vertex model we demonstrate that 3D cellular deformations drive a uniform hexatic arrangement of HCs despite large variations of the apical surface area and shape along the BP, highlighting a relationship between the relative sizes of HCs and SCs, neighbor configurations, and number ratios.

This geometric relationship implies effective repulsive forces that govern the relative displacement of HCs and SCs in the tissue. Importantly, SC-SC junctional contractility plays a pivotal role in this process by resolving HC-HC contacts. On the lateral surface, this differential apical contractility could drive the organization of HCs into a single defined stratum closer to the apical surface of BP, and SCs into multiple strata below the HCs. This quasi-stratification of HCs with a limited change in volume contributes to the expansion of HC apical surface area and hence to the hexatic arrangements. These processes collectively result in tissue-scale rearrangements within our 3D vertex model, culminating in the self-organization of HC number fractions and the emergent hexatic order observed in the developing chick BP (Fig. 4f-h).

Undergoing three-dimensional cell deformations while maintaining the cell volume, locally reduces the variation in HC apical surface area, which has recently been shown to be critical for generating hexatic order in the drosophila wing [31]. Previous studies show that oriented cell division and differential interactions between cells and with the basement membrane contribute to the vertical tissue architecture alongside the apical reorganization [32, 33]. Such differential interactions at the base are likely to contribute to the stratification of BP, since HCs lose contact with the basement membrane. In contrast, our model demonstrates that also heterotypic interactions between cell types at the apical surface contribute to stratification.

Additionally, through coupling of the lateral surface with the luminal surface, our 3D vertex model generates relative cell displacements, which result in tissue-scale flows. These cellular flows lead to a decrease in HC number density at the proximal end and an increase towards the distal end while conserving the total number of HCs in BP. Even though trans-differentiation could contribute to the distal increase of HC number, it is unlikely to contribute to the reduction of HC number in proximal parts. An alternative mechanism for generating cell flows could involve the convergent extension of the BP during development. However, this possibility is unlikely, as convergent extension would affect both HCs and SCs uniformly and cannot account for the observed relative flows between them. Future research should aim to directly measure cell flows in the developing BP to confirm and refine our understanding of the underlying mechanisms.

Variations in cellular properties along epithelial tissue axes are commonly attributed to morphogen gradients [34]. Even though our theoretical analysis shows that junctional heterogeneity is sufficient to generate the HC density and apical surface area gradients along the BP proximal-distal axis we report above, it is likely that morphogen gradients contribute to BP organization. Specifically, in developing mouse auditory epithelia [35] and hair follicles [36] they notably contribute through regulating junctional mechanics. Our work on BP provides a conceptual framework, which allows further investigation of the intricate interplay between geometrical, mechanical, and biochemical cues in three-dimensional developing tissues.

## A. Code availability

The above-described vertex model is implemented within the Utopia modeling framework for complex and adaptive systems [37]. Data handling and evaluation are performed using dantro [38]. See also www.utopia-project.org. The code of the vertex model, of the minimal lateral SC-HC-SC model and of the analysis of BP data are available on Zenodo, doi: 10.5281/zenodo.14937194, 10.5281/zenodo.14937643, and 10.5281/zenodo.14938043, respectively.

## ACKNOWLEDGMENTS

This work was supported by the Department of Atomic Energy, Government of India, Project Identification No. RTI 4006, and grants from SERB (CRG/2018/001235), TIFR Infosys-Leading Edge Grant, and Royal National Institute for Deaf People (G87) for this research. A.P. acknowledges support from International foundation for research and education via a Simons-Ashoka ECF fellowship to AP, for this research. M.R. acknowledges support from the Simons Foundation (Grant No. 287975) and DST (India) for a JC Bose Fellowship (JCB/2018/00030). J. W. thanks the University of Geneva Doc mobility programme for financial support. We thank CPDO and TI, Hasserghatta, Bengaluru for providing eggs and Central Imaging and Flow cytometry Facility (CIFF) at NCBS for microscopy imaging. Computations of the vertex model were performed at the University of Geneva’s Baobab HPC cluster. We thank Guy Richardson (Sussex Uni, Brighton, UK) for the gifts of antibodies. The Myo7a antibody developed by Dana J. Orten was obtained from the Developmental Studies Hybridoma Bank, created by the NICHD of the NIH and maintained at The University of Iowa, Department of Biology, Iowa City, IA 52242. We are grateful to the members of our laboratories for their feedback.

## Appendix A: Supplementary Figures

**FIG. S1.**
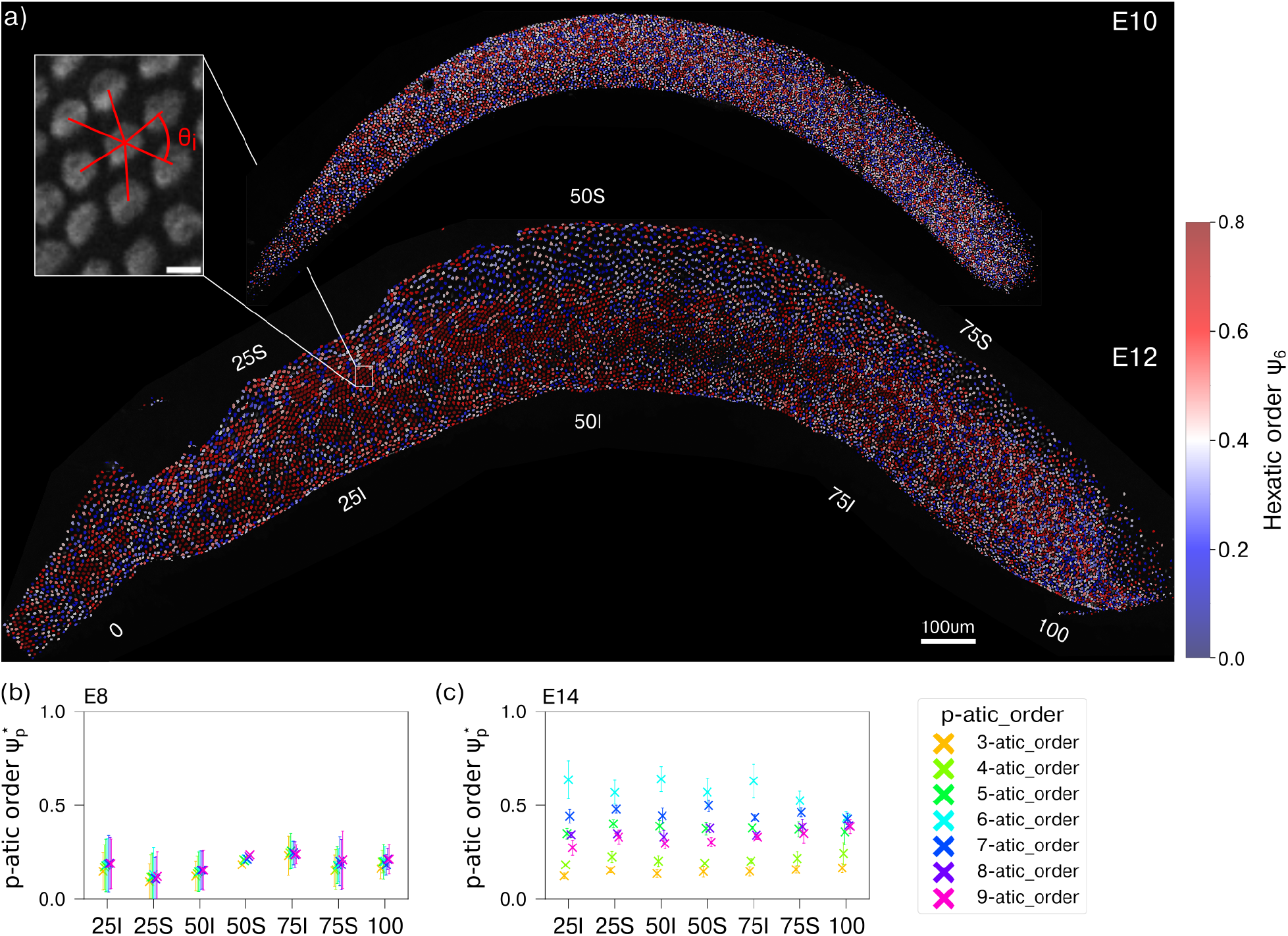
a) Hexatic order parameter *ψ*_6_ of HCs segmented in BP stained for HC antigen at E12 and E8 (lower inset). Labels indicate the P to D positions (0 % to 100 %) and S or I positions. Scale bar, 50 *µ*m. Inset shows the HCA staining, scale bar 5 *µ*m, and a sketch of angles *θ*_i_ between HC geometric centres, defining the hexatic order parameter 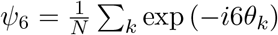 b) Indicated positions of BP at E8 stained for F-actin (green) and hair cell (HC) antigen. Scale bar, 5 *µ*m. c-d) Measurement of (c) cell area and (d) hexatic order at different positions along the P-D axis in the S and inferior (I) part of BP at stage E8 for HCs (red) and SCs (grey); see Fig. 1 for stage E14. Crosses, mean of means; errorbar, std. dev. of means.

**FIG. S2.**
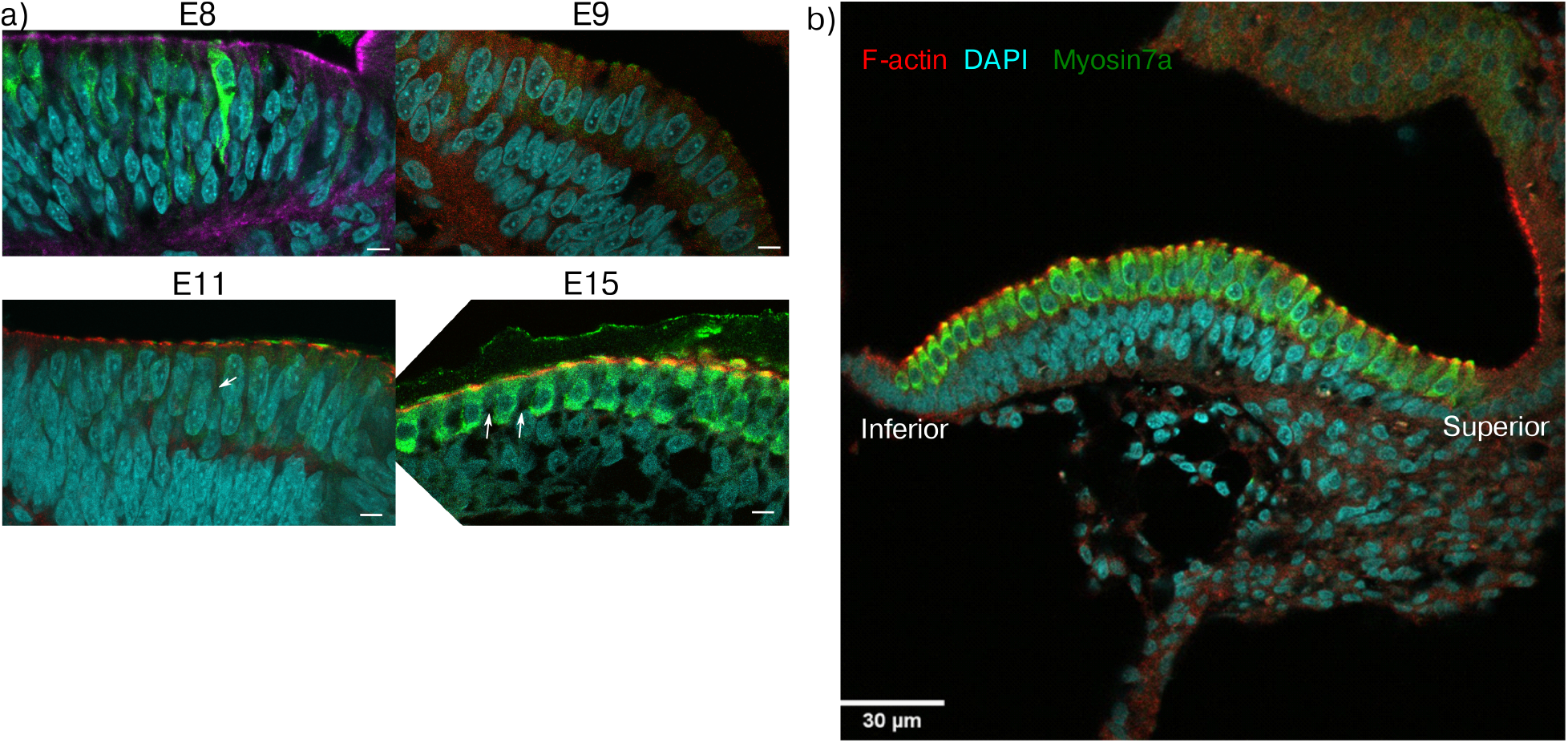
a) Cross-sections of the BP at stages E8 to E14 stained with F-actin (red), DAPI (cyan) and Myosin 7a (green).

**FIG. S3.**
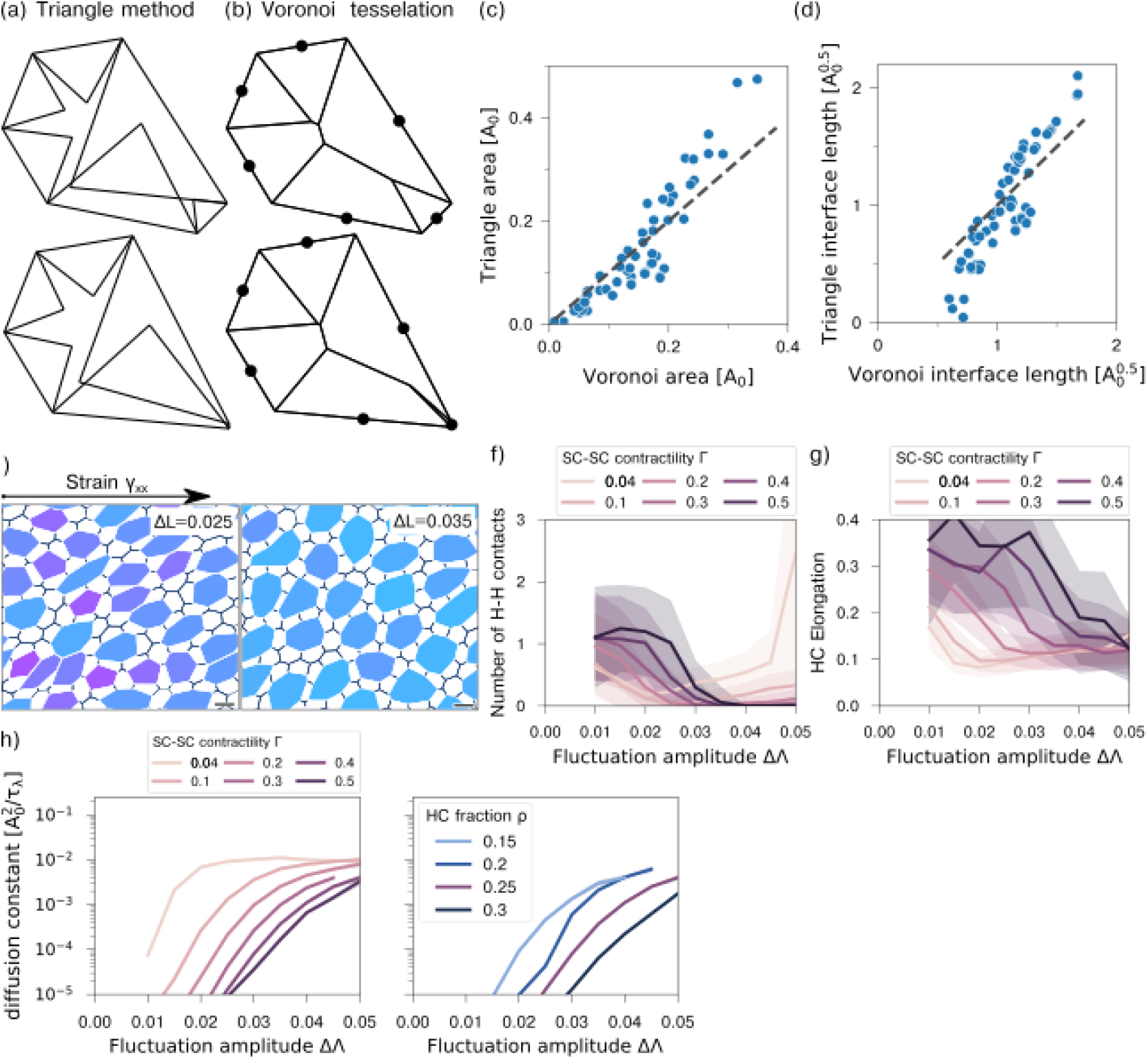
Constraints and rheology of 3D vertex model. a) Sketch of interfaces at the base of a HC in our triangle approximation. b) Voronoi tesselation using junction mid-points of the same HC base. The lower cell is undergoing a T1 transition. c-d) Comparison of sub-domain areas (c) and interface lengths (d) obtained for the Voronoi tesselation in (b) and in the triangle approximation in (a). Note, that Voronoi interface lengths are non-continuous upon T1 transitions at short junctional lengths. Such a pending transition is shown in lower configurations of (a) and (b). e) 3D vertex model while applying a continuous shear rate *v*_xx_ = 0.005*τ*_Λ_ with a low and medium linetension fluctuation amplitude ΔΛ. f-g) Measurement of (f) HC-HC contact number and (g) HC elongation at dynamic steady state undergoing continuous shear as a function of linetension fluctuation amplitude and SC-SC junctional contractility Γ. h) Measurement of diffusion constant in 3D vertex model simulation as a function of linetension fluctuation amplitude and SC-SC junctional contractility Γ.

**FIG. S4.**
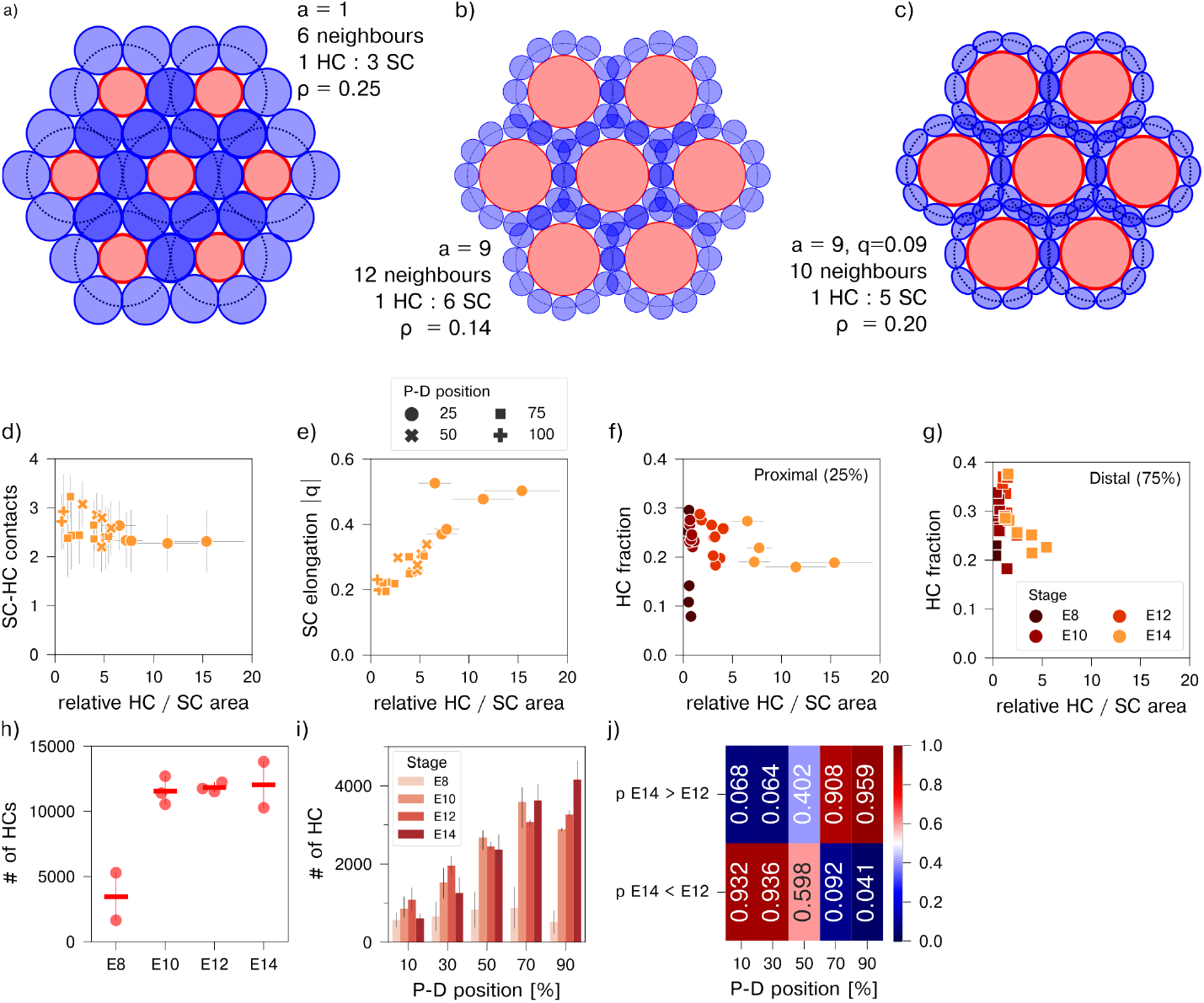
a-c) Sketch of possible HC-SC lattice structures with densely packed circular unit cells (dashed lines). Unit cells contain a single circular HC (red) and a surrounding layer of SCs (grey). From (a) to (b) the relative area of HC compared to SC was increased. The decreased diameter of SCs in (b) permits more SCs to be placed around the HC and HC fraction reduces from 0.25 to 0.14. In (c) SCs are considered ellipses with |*q*| = 0.09, reducing the number of SCs compared to at a constant area ratio. The HC fraction increased to 0.20. d-e) Mean of (d) number of SC-HC contacts and (e) SC elongation plotted against relative HC to SC area with data from BP tissues from different positions (symbols). Datapoints and crosses indicate sample means and std. dev. f-g) HC fraction plotted against relative HC to SC area with data from BP tissues from (f) 25 % and (g) 75 % at stages E8 to E14 (colour). h) HC number obtained from segmentation of whole BP HCA staining at stages E8 to E14. Dots represent individual samples, bar and errorbar represent mean and std. dev. of samples. i) HC number in 20 % intervals along the P-D axis at stages E8 to E14 (colors). Errorbars represent std. dev. of samples. j) Probability from independent T-tests whether E14 BP tissues contain more (1st row) or less (2nd row) HCs than the E12 tissues at different P-D positions. Proximal positions, 0%-40%, have less HCs at E14 than at E12 with a significance of *p <* 0.07, while distal positions, 80%-100%, have more HCs at E14 than at E12 with a significance of *p <* 0.05.

**FIG. S5.**
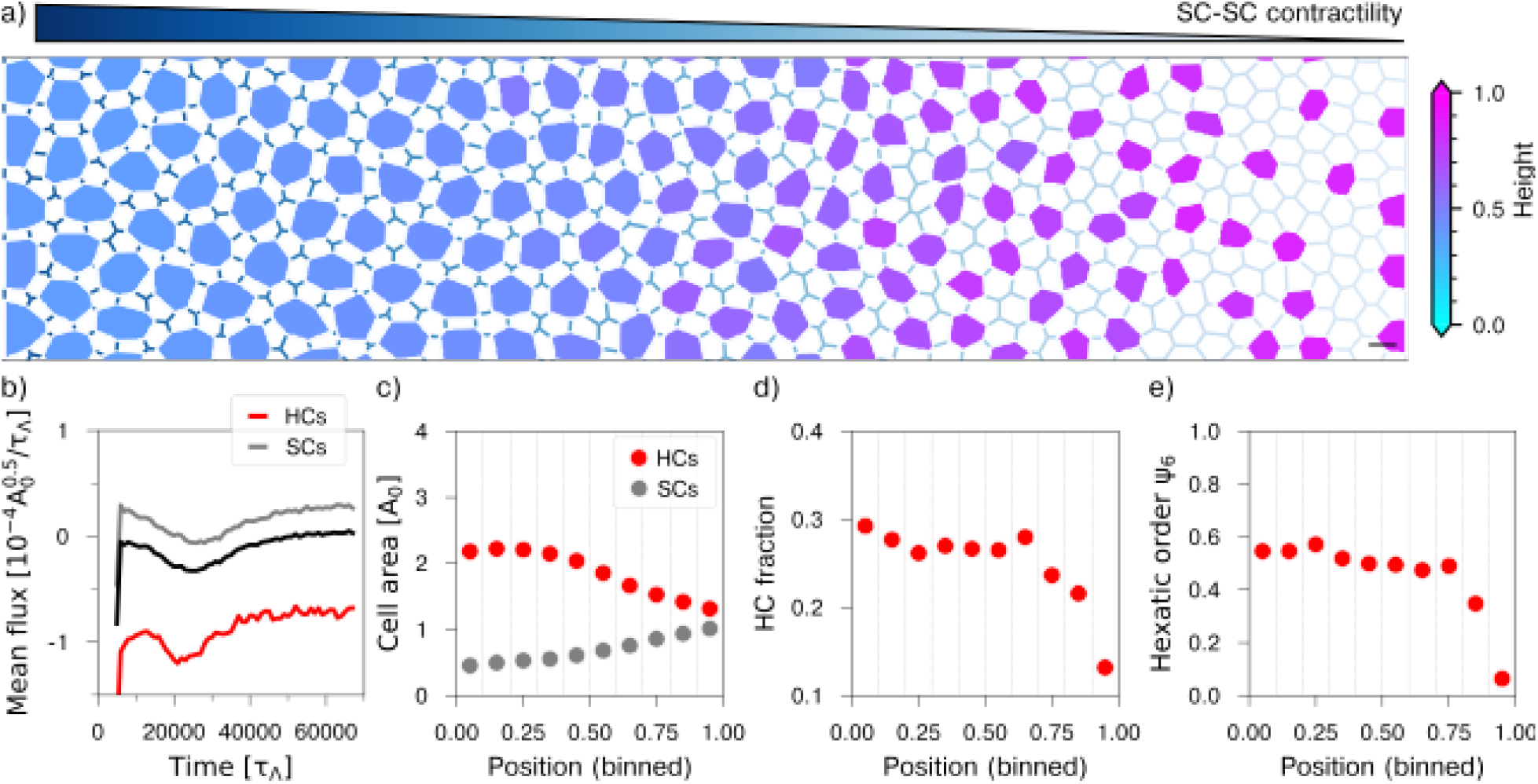
Proximal-distal gradient in apical SC-SC contractility drives cell flows. a) 3d Vertex-model simulation with graded apical SC-SC contractility. b) Mean cell fluxes for HCs (red) and SCs (gray), oriented towards the proximal and distant end, respectively, opposite to the flux in Fig. 4. Fluxes have been calculated over 5000 *τ*_Λ_ intervals. c-e) HC and SC area (c), HC fraction (d), and hexatic order (e) along the horizontal axis of the simulation at time 70 000 *τ*_Λ_.

## Appendix B: Supplementary Movies

- *Movie 1:* A movie representing junctions by Beta catenin marked in red and pp-RLC marked in green to show pp-RLC on SC-SC junctions at the apical plane.
- *Movie 2:* A 3D vertex-model simulation with graded HC extrusion speed 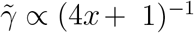 along the P-D axis position *x*. Colored cells are HCs, where color represents their

## Appendix C: Supplementary Information

### 1. Methods

#### a. Whole mount Immunostaining

Eggs, purchased from Central Poultry Development Organisation and Training Institute (Hesaraghatta, Bangalore, India), were incubated horizontally at 37 ^°^C. At the desired stage, inner ear was dissected in ice-cold PBS (without Ca2+ and Mg2+) as previously described [39]. They were fixed in 4 % PFA (dissolved in PBS) for overnight at 4 ^°^C for HCA (Guy richardson, 1:1000) and Myosin 7a (DSHB 138-1, 1:100) and 2% TCA for 20 mins at 4 ^°^C on a shaker for pp-RLC (CST 3674, 1:100) and Beta Catenin (BD Biosciences 610154, 1:200).The fixed basilar papilla was micro-dissected in PBS, permeabilised in 0.3% PBST (Tween20) for 20 mins at RT, blocked in blocking buffer (10% Goat serum 1% BSA in 0.3% PBST) at RT for 2 hours. BP was then incubated in primary antibodies diluted in blocking solution for overnight at 4 gC shaker in a 48 well plate. The samples were further washed thoroughly using PBST and further incubated in secondary antibodies (1:500), Alexa-Fluor conjugated Phalloidin (1:600) diluted in PBST for 1 hour at RT. After washing, samples were counter-stained for DAPI and then mounted in Fluoroshield.

#### b. Cross-section Immunostaining

We equilibrated fixed cochlea in 10, 20 and 30% sucrose solution before transferring to Tissue freezing medium at -25 degree. Using a cryo-microtome we sectioned the BP and obtained 20um sections on positively charged glass slides. Slides were incubated at 37^°^ C for 1 hour. Sections were them permeablised using 0.3% PBST for 30mins at room temprature and then blocked using blocking buffer (10% Goat serum 1% BSA in 0.3% PBST) at RT for 2 hours. Sections were then incubated with primary antibodies overnight at 4^°^ C. Sections are then washed thoroughly using 0.3 %PBST and incubated with secondary antibodies and Phalloidin for 1 hour at room temprature. Sections are then washed with 0.3% PBST and mounted using mounting media.

#### c. ML7 Inhibition

We used a ex-vivo organ culture method described in [39]. Briefly, we made collagen droplets from mixture containing 400ul ofrat tail collagen, 30ul of 7.5% sodium bi carbonate, 50ul of 10X DMEM and 10 ul of HEPES. We put a dissected E10 cochlea in each droplet and incubated at 37^°^ C for 10 minutes. We then added culture media (1X DMEM and N2) with DSMO or ML7 (25um) and cultured Cochlea for 4 hour in incubator maintaining 37^°^ C and 5% Co2.

#### d. Imaging

All confocal images were taken using Olympus FV 3000 inverted microscope at CIFF NCBS. Otherwise, specified images were taken using 60X oil immersion objective of numerical aperture (NA) 1.42 with step size of 0.5 *µ*m, line sequential scanning, HV values ranging between 400-500V. Whole BP images were taken using air 10X objective of NA 0.4 with step size of1 *µ*m, line sequential scanning, HV values ranging between 400-500V. It was then tiled using Fiji and Inkscape. All images are taken with 1x gain and 16bit-depth using the Olympus fluoview software. The rotation feature was used to align tissue wherever necessary.

### 2. Image Analysis

#### a. Definition of the hexatic order parameter

The hexatic order parameter quantifies the angular order of proximate HCs. For every HC it can be defined as 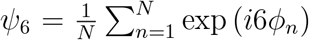, where *ϕ*_n_ is the angle of the vector from the HC’s geometric center to the center of a next-neighbor HC to the P-D tissue axis. We correct the positions of proximate HCs to the cellular elongation of the considered HC [4]. A hexatic arrangement of HCs elongated by any single axis has hexatic order *ψ*_6_ = 1. We define *ψ*_6_ = 0 for *N <* 3. More generally, we can define angular order at p-fold symmetries as 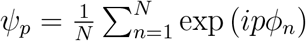, with hexatic order for *p* = 6. For HCs segmented from antigen staining, we calculate *ψ*_6_ over the neighbors in the Voronoi tesselation of the geometric centers of HCs.

#### b. Segmentation of HCs in whole BP images

Images were stitched to a whole BP image using the Stitching plugin [40] for Fiji [41]. We used the scikit-image [42] watershed algorithm to segment individual HCs from Otsu-thresholded images (Fig. 1a). Note that at late stages, E12 and E14, the cuticular plate are most intensely stained by HC antigen. Segmentation obtained in whole BP images is thus not equivalent to cell boundary segmentation in Fig. 1b. To calculate hexatic order, we performed a Voronoi tesselation of HC geometric centers and proceeded with calculation of *ψ*_6_ based on Voronoi neighborhood. The transition from an unordered to ordered hexatic phase was defined at *ψ*_6_ = 0.4 (Fig. S1a) according to data published in Prakash *et al*. [5]. All analyzed tissues from E12 and E14 BP have *ψ*_6_ *>* 0.4, in contrast, earlier stage tissues typically display lower hexatic order. Positions along the proximal-distal axis have been assigned according to the azimuthal position of the HC’s geometric centers relative to a circle segment fitted to the ensemble of HC centers.

#### c. Segmentation of cells in apical plane

For 2D morphological characterization of the tissue, we segmented cell boundaries in images obtained from confocal microscopy. To this end, we used a combination of Tissue Analyzer [43] and EPySeg [44]. Manual corrections were applied wherever necessary. We then assigned cell types based upon HC antigen staining and generated a database of cells and bonds using Tissue Analyzer. We used custom-made scripts to further quantify various parameters.

#### d. 3D HC segmentation

We manually segmented individual and sufficiently distant HCs on 2D slices. 2D slices were then combined to 3D stacks requiring zero overlap of two HCs in successive z-stacks. 3D analysis was performed using scikit-image [42].

### 3. Numerical details of the 3D vertex model

We describe a monolayer of epithelial cells using the framework of the 3D vertex model. In the vertex model, each cell is described by a polyhedron (Fig. 3a). We restrict the positions of vertices in the third dimension to approximate the specific geometry observed in the BP. SCs without contact with HCs are strictly columnar where the positions of the basal vertices are a vertical projection of the apical vertices. The SC height corresponds to the tissue height, *H*_α_ = *H*^(0)^. Similarly, HCs are also strictly columnar. However, their height is an independent variable of the system, 0 ≤ *H*_α_ ≤ *H*^(0)^. Thus, the basal vertices of HCs have in general at different z-position than those of SCs. Yet, their x-y-position remains a projection of the apical plane. The interfaces and volumes below a HC are attributed to the neighbouring SCs in a triangle approximation. These SCs thus deviate from a columnar geometry.

#### a. Geometric restrictions in 3D

In the triangle approximation, the HC’s basal surface is divided into *n*_S_ isosceles triangles, one for each SC neighbour of the HC (Fig. S3a). The base of each triangle has length *l*_α,β_ and coincides with the junction shared by the HC and SC. The length of the isosceles edges is proportional to the length of the base, such that the surface of the triangle is given as

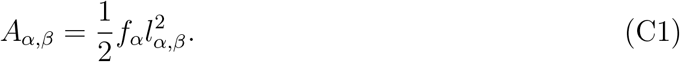

The proportionality factor *f*_α_ is chosen such that the total triangle surfaces is equal to the area of the HC’s base, *A*_α_ = ∑_β_ *A*_α,β_.

The volume of HCs and SCs can thus be explicitly written as

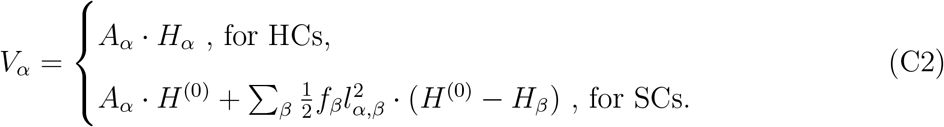

Similarly, we can write the cells’ surface area as

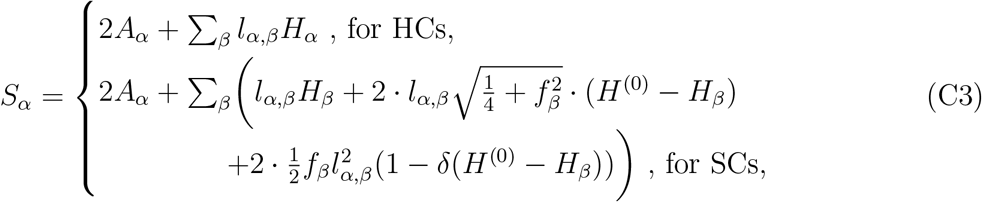

where *δ*(*H*^(0)^ − *H*_β_) = 1 for *H*^(0)^ = *H*_β_ and 0 otherwise. Note, that in this approximation the identification of an interface between neighbouring cells is not possible. For instance, two SCs separated by a HC might share an interface in the Voronoi tesselation of the same HC (central lines in Fig. S3b). However, when assuming the same parameter values for all interface types this distinction is not necessary. Note that in BP heterogeneities between interface types, originating from pp-RLC localisation, are located at the apical side of the tissue, where interfaces are well-defined.

To compare our geometric simplification to an accurate representation of volumes and interfaces, we measured the triangle areas and interfaces, as well as their Voronoi tesselation equivalents, in various geometric arrangements (Fig. S3c,d). We define the interfaces as the perimeter minus the junction’s length, thus considering only the interface shared with other cells. We observe that both, the triangle areas and interfaces, correlate with the Voronoi tesselation, Pearson Correlation Coefficient r = 0.93 and r = 0.90 for area and interfaces respectively. Note, that only the triangle interface length vanishes for very short junctions, while Voronoi interfaces remain finite for short junctions.

#### b. Minimization of the work function

We consider the polyhedron arrangement at the mechanic equilibrium, obtained using the steepest-descent method. In each step, we allow for changes in the work function (eq. 2) caused by remodeling of the cellular structure defined by the junctions. Explicitly, whenever the length of an apical cell-cell junction is shorter than a threshold length *l*_min_, a so-called T1 transition occurs. In this transition, the junction is removed and a new junction of length *l* = *d* · *l*_min_ is created in orthogonal orientation [25]. We discard transitions that increase the work function. Likewise, the removal of cells with below threshold apical area or volume could be considered, often referred to as a T2 transition. However, in the BP no cell apoptosis is observed. To prevent T2 transitions in our vertex-model, we consider an elastic contribution in our work-function for cells below a minimum apical area, 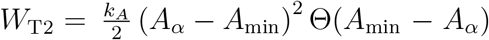, with the Heavidside step function Θ(*A*_min_ − *A*_α_) = 1 for *A*_α_ ≤ *A*_min_ = 0.2 and 0 otherwise.

Remodeling of the cellular structure permits a viscous relaxation of the tissue after persistent deformation and occurs on a long timescale. Fast deformations typically result in an elastic response without remodeling of the cellular structure [29, 45]. To obtain smooth stress-strain curves, we relax the accumulated strain by introducing fluctuations in the system. We follow Duclut *et al*. [29] and adopt fluctuations in the bond tension parameter, where the dynamics of individual bonds are described by an Ornstein-Uhlenbeck process,

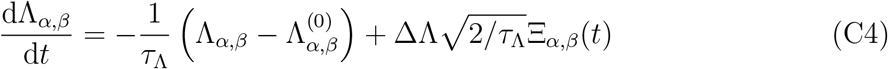

Here the first term on the right-hand side describes a relaxation of the parameter to a mean value Λ_α,β_ over a timescale given by the characteristic time *τ*_Λ_. The second term comprises the Gaussian white noise Ξ_α,β_(*t*) with zero mean ⟨Ξ_α,β_(*t*)⟩ = 0 and correlations ⟨Ξ_α,β_(*t*)Ξ_µ,ν_(*t*^′^)⟩ = *δ*_α,β_*δ*_µ,ν_*δ*(*t*−*t*^′^), with *δ*_α,β_, *δ*_µ,ν_*δ*(*t*−*t*^′^) = 1 when {*α, β*} and {*µ, ν*} describe the same junction and *t* = *t*^′^ and 0 otherwise. The magnitude of the fluctuations is captured by the parameter ΔΛ. We have chosen the characteristic time scale of the fluctuations *τ*_Λ_ to be much larger than the elastic relaxation time of the work function.

The initial state of our model is generated from an arrangement of *N*_x_×*N*_y_ hexagonal cells in a periodic domain with area *A*_domain_ = *NV* ^(0)^*/H*^(0)^ such that the cells fit into the domain with no volume constraint. We subsequently minimise the work function over multiple characteristic times of the fluctuations, providing sufficient time for T1 transitions to occur. For determining the rheology of our vertex model, we use Lees-Edwards boundary conditions, where the upper and lower domain replicas move at a constant velocity *u*_x_ = ±*ν* [46].

For simulations with gradients along the horizontal axis, we consider fixed boundary conditions to the left and right end of the tissue, where boundary vertices are free to move along the y-axis but are placed within a quadratic potential in the x-axis. The distance of the left and right boundary is *L*_x_ = *N*_x_ × *w*, with *w* the width of a hexagon of target volume and height. At the top and bottom, we consider periodic boundary conditions.

#### c. Vertex model simulation with graded junctional contractility

In the main text, we discuss a spatial gradient in HC area growth speed and vary the associated dissipation rate γ̄, see Fig. 4. Alternatively, this difference in HC area growth speed and gradient in HC apical area by E14 could result from graded apical SC-SC contractility, compare Fig. 3c,e. We explore this possibility by setting 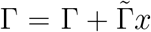, with *x* the normalized geometric center of a junction, *x* ∈ [0, 1] (Fig. S5a).

We observe that also in this situation proximal HCs grow faster in apical area due to the stronger SC-SC apical contractility (Movie X). Furthermore, we observe a flux of HCs against the gradient in SC-SC contractility towards the proximal side (Movie X, Fig. S5b). This is opposite to the flux observed when applying a gradient in HC extrusion rate. With graded SC-SC contractility, the flux is independent of HC area growth and 3D remodeling, as the flux is maintained even when height remodeling is turned off, γ̄ = 0. Instead, this flux results from the heterotypic surface interactions, which also drive HC-HC separation. Even though the removal of a HC-HC interface during a T1 transition results in the addition of a SC-SC interface, it still reduces the value of the work function through the rearrangement of multiple cells. Indeed, the HC-SC interface can be maximized in the absence of HC-HC contacts, such that the SC-SC interface length is minimized. The increased contractility of SC-SC junctions toward the proximal side drives the HC flux that permits to further increase HC-SC interface length and decrease SC-SC interface length in proximal parts.

Similar to our results in Fig. 3, at steady state, HC apical area is largest in proximal positions where SC-SC contractility is largest and decreases along the gradient of SC-SC contractility (Fig. S5c). Furthermore, as a result of HC flux, HC fraction increases proximally (Fig. S5d) and effective HC-HC repulsion hinders further HC area growth in proximal positions (Fig. S5c,d) compared to results in Figure 4. Throughout most of the tissue we observe hexatic order (Fig. S5e) similar to previous results. In contrast, in the distal part of the tissue, HCs become scarce and, due to low SC-SC contractility, HC area remains small. This results in low hexatic order.

Since the HC fraction does not agree with our observations in BP, we consider this scenario to play a less prominent role in BP development than a gradient in cell extrusion rate.

### 4. Analytical estimation of HC fraction

In the following, we give an analytical estimate of the HC fraction. We consider a regular arrangement, where any two HCs are separated by a single layer of SCs. Specifically, we consider a 2D array of circular unit cells forming a hexagonal lattice (Fig. S4a,b). We divide every unit cell of radius *R* into a single central HC of radius *r*_h_ and *N* neighboring SCs of radius *r*_s_, with *R* = *r*_h_ + *r*_s_. The HC fraction *ρ* is then given by

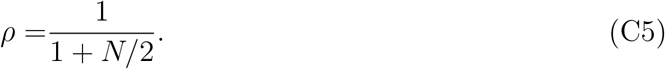

In BP, we find that SCs contact 2 to 3 HCs (Fig. S4d) justifying the use of *N/*2 in the expression for *ρ*.

The maximum number *N* of SCs surrounding each HC can be estimated as

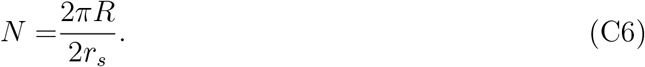

To compare this to our data from BP samples from different positions and stages and numerical data in Fig. 4e, we express *N* in terms of the relative area *a* of HCs to SCs,

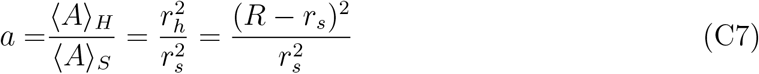

such that

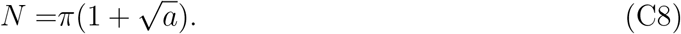

We observe that the first scenario this estimate provides an upper limit to our experimental and numerical data (Fig. 4f). However, BP tissues often show significantly lower neighbor numbers. In the BP, SC apices are often not as circular as HCs but are squeezed between two HCs and thus elongated (Fig. 1b). We account for SC elongation *q* and describe SC as ellipses with long and short axes *r*_s_ exp (2*q*) and *r*_s_ exp (−2*q*), respectively. Aligning the SC long axes with the HC tangents (Fig. S4c), we obtain

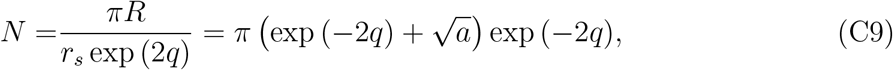

with *R* = *r*_s_ exp (−2*q*) + *r*_h_. We fit the SC elongation to the data from BP at E14 in Figure 4f, obtaining *q* = 0.129(14). In BP, we measure slightly larger *q* values (Fig. S4e). Our analytical estimate Eq. C5 recapitulates well the HC fraction obtained from individual images of different BP positions at E14 (Fig. 4g). We also find a good agreement between our estimate and our simulated tissues regarding HC neighbour number and HC fraction (Fig. 4f,g). Furthermore, hexatic order in simulated BP is similar to E14 BP (Fig. 4h).

## Appendix D: Supplementary tables

**TABLE I.** Typical parameter values used in the 3D vertex model.

| Symbol | Description | Value | Unit |
| --- | --- | --- | --- |
| $K_V, K_S$ | Elastic constants | 1 | 1, $K_V(V^{(0)})^2$ |
| $V_\alpha^{(0)}$ | Target cell volume | 1 | 1 |
| $s_\alpha^{(0)}$ | Target cell shape index | 5.78 | 1 |
| $H^{(0)}$ | Target tissue height | 1 | $(V^{(0)})^{1/3}$ |
| $\Lambda_{\alpha,\beta}$ | Surface tension | 0 | $K_V(V^{(0)})^{4/3}$ |
| $\Gamma_{\alpha,\beta}$ | Active apical junction contractility | 0 – 0.6 | $K_V(V^{(0)})^{4/3}$ |
| $\tilde{\Gamma}_{\alpha,\beta}$ | Active apical perimeter contractility | 0.01 | $K_V(V^{(0)})^{4/3}$ |
| $\Delta\Lambda$ | Apical line tension fluctuations magnitude | 0.01-0.05 | $K_V(V^{(0)})^{5/3}$ |
| $N$ | Total number of cells | 512 | |
| $\rho_H$ | Hair cell number density | 0.1-0.4 | |
| $\gamma$ | Time step for numerical integration | 0.05 | a.u. |
| $\tau_{\text{relax}}$ | Elastic relaxation time | 160 | numerical steps |
| $\tau_\Lambda$ | Characteristic time of bond fluctuations | 3 | $\tau_{\text{relax}}$ |
| | Simulated time after cell divisions | 240 | $\tau_\Lambda$ |
| $l_{\min}$ | Threshold length of a junction for T1 remodeling | 0.03 | $(V^{(0)})^{1/3}$ |
| $d$ | Separation factor for new junction in T1 remodeling | 1.05 | |
| $A_{\min}$ | Minimum apical area | 0.2 | $(V^{(0)})^{2/3}$ |
|  | Number of simulations per parameter set | 8 |  |

**TABLE II.** Parameter values used in the simulation of HC extrusion with increasing SC-SC contractility Γ (Fig. 3c,e). Other parameters as in table I.

| Symbol | Description | Value | Unit |
| --- | --- | --- | --- |
| $\tilde{\gamma}$ | Dissipation rate of HC height change | 2.E-6 | dt |
| $\Gamma_{S,S}$ | SC-SC junctional contractility | 0.4 | $K_V(V^{(0)})^{4/3}$ |
| $\Delta\Lambda$ | Apical line tension fluctuations magnitude | 0.035 | $K_V(V^{(0)})^{5/3}$ |
| $\rho_H$ | Hair cell number density | 0.25 | |
| $T_{\max}$ | Total simulated time | 50 000 | $\tau_\Lambda$ |

**TABLE III.** Parameter values used in the simulation of HC extrusion (Fig. 3d,f,g). Other parameters as in table I.

| Symbol | Description | Value | Unit |
| --- | --- | --- | --- |
| $\tilde{\gamma}$ | Dissipation rate of HC height change | 1. | dt |
| $\Delta\Gamma_{S,S}$ | Incremental of SC-SC junctional contractility | 1.E-5 | $K_V(V^{(0)})^{4/3}$ |
| $\Delta\Lambda$ | Apical line tension fluctuations magnitude | 0.03 | $K_V(V^{(0)})^{5/3}$ |
| $T_{\max}$ | Total simulated time | 50 000 | $\tau_\Lambda$ |

**TABLE IV.**
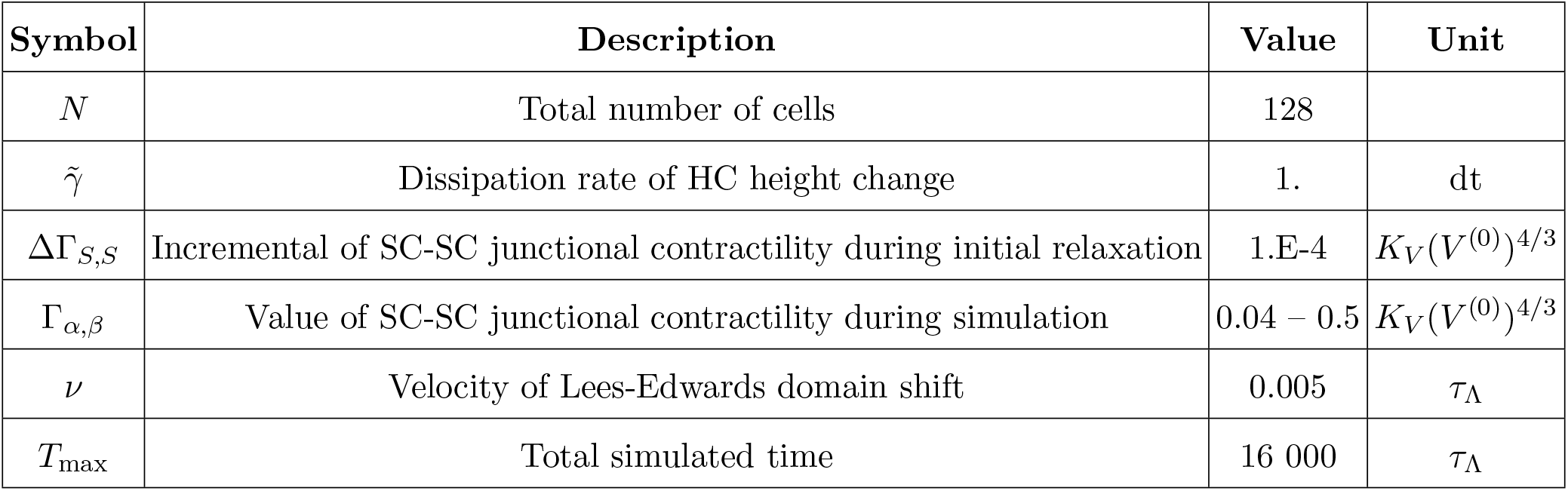
Parameter values used in the rheological test (Fig. S3e-g). Other parameters as in table I.

| Symbol | Description | Value | Unit |
| --- | --- | --- | --- |
| $N$ | Total number of cells | 128 | |
| $\tilde{\gamma}$ | Dissipation rate of HC height change | 1. | dt |
| $\Delta\Gamma_{S,S}$ | Incremental of SC-SC junctional contractility during initial relaxation | 1.E-4 | $K_V(V^{(0)})^{4/3}$ |
| $\Gamma_{\alpha,\beta}$ | Value of SC-SC junctional contractility during simulation | 0.04 – 0.5 | $K_V(V^{(0)})^{4/3}$ |
| $\nu$ | Velocity of Lees-Edwards domain shift | 0.005 | $\tau_\Lambda$ |
| $T_{\max}$ | Total simulated time | 16 000 | $\tau_\Lambda$ |

**TABLE V.**
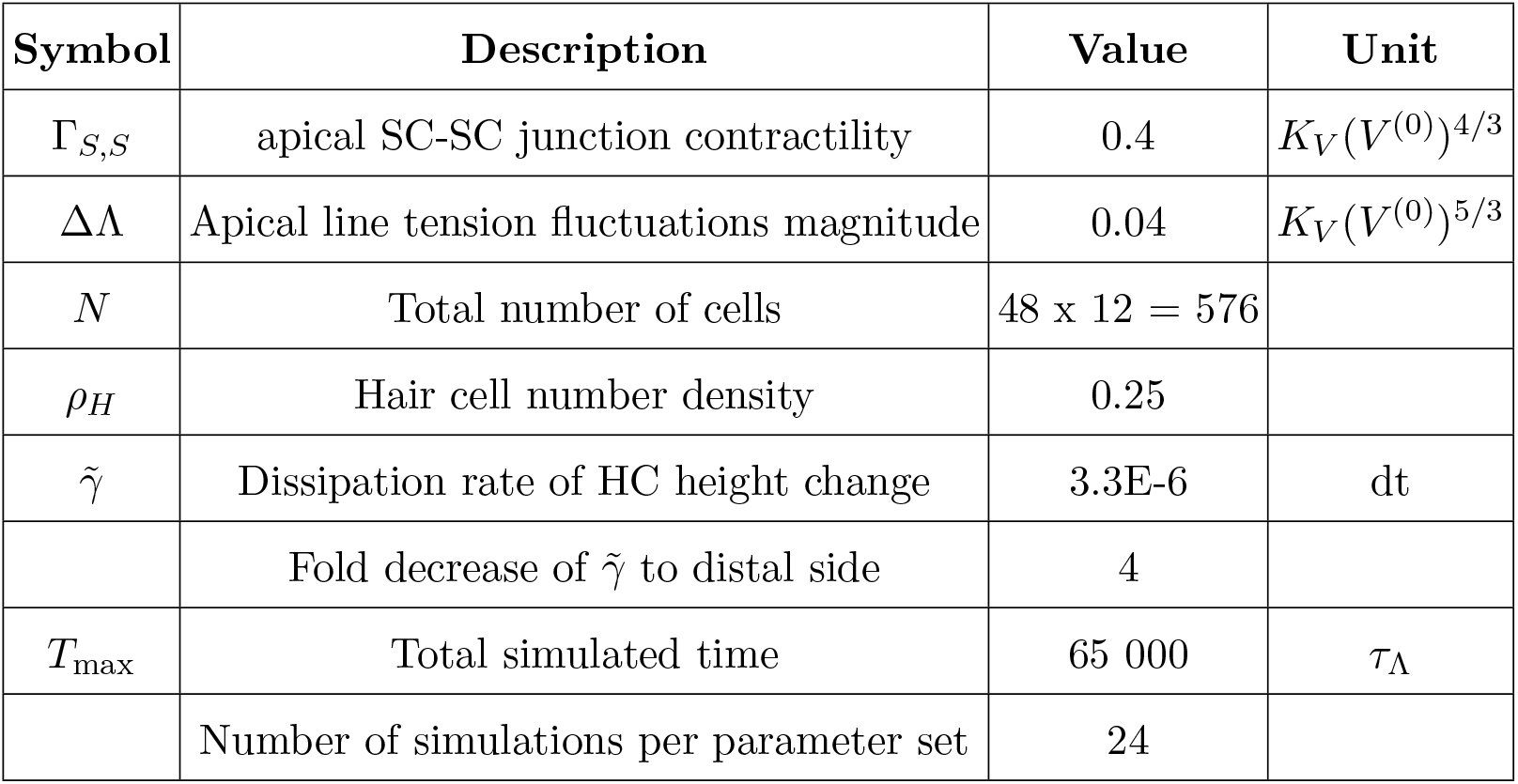
Parameter values used in simulations with graded HC extrusion dissipation rate (Fig. 4). Other parameters as in Table I.

**TABLE VI.** Parameter values used in simulations with graded SC-SC contractility (Fig. S5). Other parameters as in Table I.

| Symbol | Description | Value | Unit |
| --- | --- | --- | --- |
| $\Gamma_{S,S}$ | Proximal apical SC-SC junction contractility | 0.35 | $K_V(V^{(0)})^{4/3}$ |
| $\Gamma_{S,S}$ | Distal apical SC-SC junction contractility | 0.05 | $K_V(V^{(0)})^{4/3}$ |
| $\Delta\Gamma_{S,S}$ | SC-SC junction contractility gradient | -0.3 | $K_V(V^{(0)})^{4/3}$ |
| $\Delta\Lambda$ | Apical line tension fluctuations magnitude | 0.04 | $K_V(V^{(0)})^{5/3}$ |
| $N$ | Total number of cells | 48 x 12 = 576 | |
| $\rho_H$ | Hair cell number density | 0.25 | |
| $\tilde{\gamma}$ | Dissipation rate of HC height change | 3.3E-6 | dt |
| $T_{\max}$ | Total simulated time | 65 000 | $\tau_\Lambda$ |
|  | Number of simulations per parameter set | 16 |  |

